# TGFβ primes alveolar-like macrophages to induce type I IFN following TLR2 activation

**DOI:** 10.1101/2024.09.04.611226

**Authors:** Sean M. Thomas, Abigail P. McGee, Taryn E. Vielma, Laurisa M. Ankley, Alexander W. Rapp, Kayla N. Conner, Francois LeSage, Christopher D. Tanner, Eleanor C. Scheeres, Joshua J. Obar, Andrew J. Olive

**Author notes:** Corresponding authors: Andrew Olive, Michigan State University, College of Osteopathic Medicine, Department of Microbiology, Genetics, and Immunology, East Lansing, MI 48824, E-mail address, Joshua J. Obar, Geisel School of Medicine at Dartmouth, Department of Microbiology & Immunology, 1 Medical Center Drive, Lebanon, NH 03756. These authors contributed equally (Abigail P. McGee, Taryn E. Vielma, Laurisa M. Ankley).

## Abstract

Alveolar macrophages (AMs) are key mediators of lung function and are potential targets for therapies during respiratory infections. TGFβ is an important regulator of AM differentiation and maintenance, but how TGFβ directly modulates the innate immune responses of AMs remains unclear. This shortcoming prevents effective targeting of AMs to improve lung function in health and disease. Here we leveraged an optimized *ex vivo* AM model system, fetal-liver derived alveolar-like macrophages (FLAMs), to dissect the role of TGFβ in AMs. Using transcriptional analysis, we first globally defined how TGFβ regulates gene expression of resting FLAMs. We found that TGFβ maintains the baseline metabolic state of AMs by driving lipid metabolism through oxidative phosphorylation and restricting inflammation. To better understand inflammatory regulation in FLAMs, we next directly tested how TGFβ alters the response to TLR2 agonists. While both TGFβ (+) and TGFβ (-) FLAMs robustly responded to TLR2 agonists, we found an unexpected activation of type I interferon (IFN) responses in FLAMs and primary AMs in a TGFβ-dependent manner. Surprisingly, mitochondrial antiviral signaling protein and the interferon regulator factors 3 and 7 were required for IFN production by TLR2 agonists and the IFN response was dependent on mitochondrial reactive oxygen species. Together, these data suggest that TGFβ modulates AM metabolic networks and innate immune signaling cascades to control inflammatory pathways in AMs.

## INTRODUCTION

The pulmonary space is a specialized environment evolved to facilitate gas exchange and maintain lung function (1, 2). To protect against exposures to airborne microorganisms and particulates, lung alveoli contain a specialized phagocyte population, alveolar macrophages (AMs) (2, 3). These AMs, like many other tissue-resident macrophages, seed the lungs from the fetal liver and serve two primary purposes: to preserve lung homeostasis by maintaining optimal surfactant levels in the lungs and to patrol the alveolar space for inhaled debris, initiating an inflammatory response when necessary (4–6). Given the importance of maintaining pulmonary function, AMs must strictly regulate their inflammatory responses to prevent unnecessary inflammation and tissue damage (7, 8). Compared to other inflammatory macrophages, including bone marrow-derived macrophages (BMDMs), AMs are more hypo-inflammatory against many pathogenic stimuli, a characteristic that is mediated by their distinct ontogeny and the lung environment (8–10). In fact, circulating monocytes that are recruited to the lungs following infection have been shown to adapt to the local environment and take on AM-like phenotypes (11). Two key cytokines, GM-CSF and TGFβ, are known to mediate AM differentiation and functions in the lung environment (6, 12–14). While the role of GM-CSF is better understood due to its importance in preventing pulmonary alveolar proteinosis, how TGFβ directly modulates the AM state and function remains unclear, limiting our ability to target AMs and improve lung function in health and disease.

TGFβ exists as three separate isoforms (TGFβ1-3) that all bind to the same TGFβ coreceptors (TGFβRI, TGFβRII) (15). TGFβ-1 is primarily produced by macrophages, but in an inactive form, conjugated with a latency-associated peptide (LAP) (16–18). Inactive TGFβ-1 (referred to as TGFβ from here on) is activated following enzymatic, acidic, or receptor-mediated cleavage of the LAP from TGFβ (18, 19). In the lungs, inactive TGFβ is primarily produced by AMs which is then activated by the alveolar epithelial type II cells (AECII) through the activity of the αvβ6 integrin on alveolar epithelial cells (6, 14, 18). Thus, maintaining AMs requires unique interactions between the lung epithelium and disruptions of this environment results in dysregulated pulmonary responses.

In its active form, TGFβ is a versatile cytokine that triggers SMAD complex translocation to the nucleus to drive a multitude of processes, including stem cell differentiation, chemotaxis, and immune regulation, depending on the context in which it is acting (20, 21). Much of this heterogeneity in cellular responses to TGFβ is thought to be due to crosstalk between other transcriptional regulators and epigenetic regulation (22). In the lungs, TGFβ plays critical roles both in lung development and disease. Mice lacking any of the three isoforms of TGFβ or either of the two receptors have varying degrees of deformed lung structure and alveologenesis due to dysregulated interactions between the lung epithelium and mesenchyme during development (23–26). TGFβ is also implicated in the development of idiopathic pulmonary fibrosis (IPF) through its induction of myofibroblast differentiation from lung fibroblasts and suppression of anti-fibrotic factors prostaglandin E2 and hepatocyte growth factor production (27–29). Given the importance of TGFβ to maintain AMs in the lungs, it is essential to better understand how TGFβ modulates the inflammatory potential of AMs.

Fully dissecting the role of TGFβ in AM regulation requires *ex vivo* models that recapitulate key aspects of the lung environment. Recent work by several groups showed that growth of macrophages in both GM-CSF and TGFβ stabilizes the AM-like state for cells grown in culture (30–32). We recently optimized the fetal liver-derived alveolar-like macrophages (FLAMs) model which propagates fetal liver cells in both GM-CSF and TGFβ, allowing for long-term propagation and genetic manipulation of cells that recapitulate many aspects of AM functions (32). Removing TGFβ from these cells results in a loss of the AM-like state, such as decreased expression of the key AM transcription factor peroxisome-proliferating activating receptor gamma (PPARγ) and increased expression of the LPS co-receptor CD14. These data suggest that TGFβ not only maintains the AM state but plays an important role in modulating the inflammatory response of AMs.

In this report we directly examine how TGFβ shapes AM function and inflammatory responses. Using transcriptional analysis, we globally defined how TGFβ regulates the gene expression of resting FLAMs, identifying a key role of TGFβ in maintaining the metabolic state of AMs. In parallel, we characterized how TGFβ shapes the inflammatory response of AMs following the activation of toll-like receptor 2 (TLR2), uncovering an unexpected link between TGFβ, TLR2, and type I interferon (IFN). We found that a range of TLR2 agonists activate type I IFN responses in a TGFβ-dependent manner. Further, mechanistic studies found this type I IFN response required the transcription factors interferon regulatory factor 3 and 7 (IRF3/7) in addition to the mitochondrial antiviral signaling adaptor (MAVS) protein. We also found increased peroxisome and mitochondrial organelle staining in response to TGFβ, resulting in increased reactive oxygen species following TLR2 activation which contributed to IFN production. Overall, these data suggest that TGFβ rewires the AMs and potentiates the activation of unique innate immune signaling not observed in other macrophage populations.

## RESULTS

### TGF**β** drives lipid metabolism, restrains cytokine expression, and maintains FLAMs in the AM-like state

We previously developed FLAMs as an *ex vivo* model of AMs to understand the mechanistic signals and regulatory networks that maintain cells in the AM-like state (32). TGFβ is a key cytokine needed to maintain AMs *in vivo* and to maintain FLAMs in the AM-like state, yet how TGFβ modulates AM functions and transcriptional networks remains poorly resolved (6, 14, 32). As a first step, we confirmed that TGFβ is required to broadly maintain the AM-like state in FLAMs. FLAMs were grown in medium containing GM-CSF with or without TGFβ for two-weeks and the expression of the AM-associated surface markers SiglecF and CD14 were quantified by flow cytometry. We observed that FLAMs grown with TGFβ had high expression of SiglecF and CD11a together with low expression of CD14 and CD11b on their surface, while FLAMs grown without TGFβ had opposite pattern of expression with low expression of SiglecF and CD11a and higher expression of CD14 and CD11b (Figure 1A). Since PPARγ is a key transcription factor in AMs and is expressed in both AMs and FLAMs (32, 33), we measured the effect of TGFβ on PPARγ expression. mRNA was isolated from FLAMs grown with and without TGFβ and quantitative RT-PCR was used to determine the expression of *Pparg*. FLAMs grown with TGFβ maintained higher expression of *Pparg*, while cells grown in the absence of TGFβ showed significantly decreased *Pparg* expression (Figure 1B). These data confirm that TGFβ helps maintain FLAMs in an AM-like state, similar to what has been seen with TGFβ’s role in differentiation and maintenance of AMs in the lungs (6, 32).

**Figure 1.**
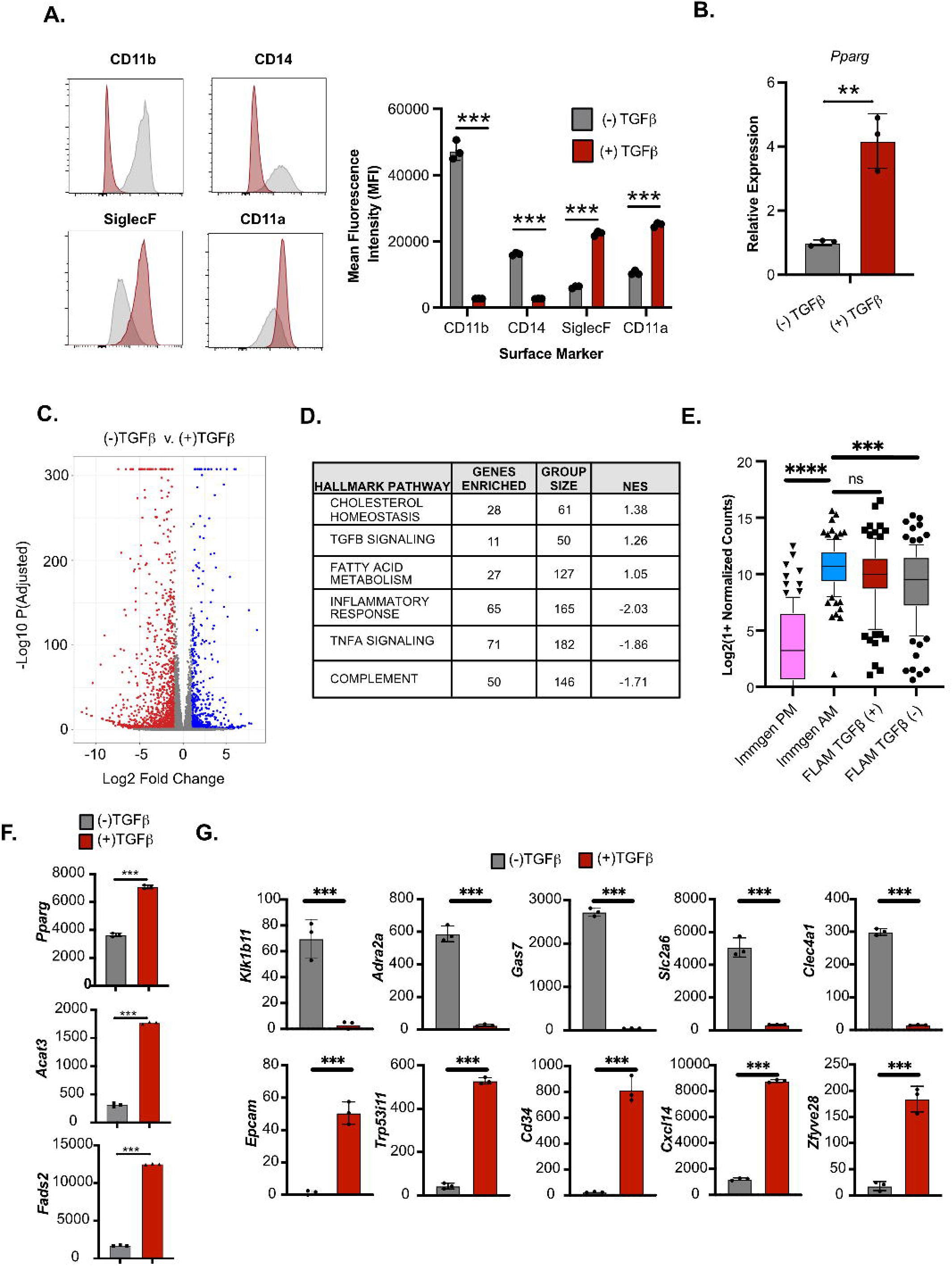
TGFβ drives lipid metabolism, restrains chemokine expression, and maintains FLAMs in the AM-like state. **(A)** The expression of the indicated surface markers was quantified by flow cytometry from TGFβ (+) and TGFβ (-) FLAMs and are shown as representative histograms (left) and the quantified mean fluorescence intensity (right). Each point represents a technical replicate from one representative experiment of 3. **p<0.01 by unpaired students t-test. **(B)** *Pparg* expression was quantified by qRT-PCR using 2^(-ΔΔCT)^ relative to GAPDH in untreated TGFβ (+) and TGFβ (-) FLAMs. Each point represents a technical replicate from one representative experiment of 3. **p<0.01 by unpaired students t-test. **(C)** Differentially expressed genes were identified between untreated TGFβ (+) and TGFβ (-) FLAMs. Red points represent significantly under-expressed genes (adjusted p-value < 0.05) and blue points represent significantly overexpressed genes (adjusted p-value < 0.05) between TGFβ (+) and TGFβ (-) FLAMs. Each point represents the mean of three biological replicates from one experiment. DESeq2 was used to determine significance using the adjusted p-value to account for multiple hypothesis testing. **(D)** Shown are the top hallmark pathways enriched in TGFβ (+) and TGFβ (-) FLAMs. **(E)** Normalized counts of core AM genes were compared among TGFβ (+) and TGFβ (-) FLAMs and previously published datasets from ImmGen (ImmGen PM and ImmGen AM; accession GSE122108, https://www.ncbi.nlm.nih.gov/geo/query/acc.cgi?acc=GSE122108). The box plot shows the median with quartiles representing the 10th to 90th percentile range of the data within that cell type. Each point represents the mean normalized counts of an individual gene. The Mann–Whitney *U* test was used to make statistical comparisons between each cell type and to compare medians. **(F)** Gene expression was quantified from normalized counts for key genes important in lipid metabolism. **(G)** Gene expression was quantified from normalized counts for a subset of genes differentially expressed from Yu et al. (6). For (F) and (G) each point represents a technical replicate from one experiment. *** adjusted p-value <0.001 using DeSeq2 analysis.

To better understand how TGFβ globally regulates FLAMs, we next conducted whole-transcriptome RNA sequencing analysis on FLAMs grown in the presence and absence of TGFβ. Differential expression analysis identified hundreds of genes that were significantly changed between FLAMs grown with or without TGFβ (Figure 1C and GSE276577 for normalized count table). To globally identify pathways that were uniquely enriched in TGFβ (+) FLAMs, we employed gene set enrichment analysis (GSEA), using a ranked gene list generated from the differential expression analysis (Figure 1D). Among the top hallmark pathways enriched in TGFβ (+) FLAMs were cholesterol homeostasis, TGFβ signaling, and fatty acid metabolism (Figure 1D and Supplemental Figure 1A). Supporting this observation, TGFβ (+) FLAMs contained significantly less intracellular lactate than TGFβ (-) FLAMs (Supplemental Figure 1B). Given that AMs are known to drive PPARγ-dependent lipid metabolism, these data suggest the FLAM transcriptional profile is similar to primary AMs (34, 35). In contrast, pathways enriched in TGFβ (-) FLAMs included the inflammatory response, TNFα signaling, and complement (Figure 1D and Supplemental Figure 1A). To directly test similarities between FLAMs and AMs, we compared the expression of a previously published AM gene signature in FLAMs grown with and without TGFβ in addition to primary AMs and peritoneal macrophages (PMs) for the immunological genome project (ImmGen) (Figure 1E and Supplemental Table 1) (36). While there was no significant difference in the expression of this gene signature between AMs and TGFβ (+) FLAMs, there was a significant difference between AMs and TGFβ (-) FLAMs supporting that TGFβ maintains FLAMs in an AM-like state. We next directly compared the expression of a subset of genes related to differentially expressed pathways (Figure 1F). We found high expression of lipid metabolism genes including *Pparg, Acat3*, and *Fads2* in TGFβ (+) FLAMs. These data support a key role of TGFβ in maintaining the AM-specific metabolic state To examine the relevance of FLAMs as a model of primary AMs, we compared our transcriptomics with those identified from Greter and colleagues who examined the transcriptome of sorted primary AMs with and without the TGFβR (6). Of the top 15 genes that Yu et al. found to be upregulated or downregulated from AMs in the lungs, 13 of 15 upregulated and 12 of 15 downregulated genes were similarly differentially expressed significantly in TGFβ (+) and TGFβ (-) FLAMs. (Figure 1G and Supplemental Table 1). These data strongly suggest that FLAMs recapitulate the biology observed in the lungs of intact animals and provide a useful tool to dissect the underlying mechanisms of TGFβ on AMs.

### TGFβ mediates a type I IFN response in AMs following Pam3CSK4 Activation

To understand how TGFβ alters the innate immune response of AMs we next directly tested the response of FLAMs grown with and without TGFβ to inflammatory stimuli. Many bacterial and fungal respiratory infections activate TLR2 signaling during infection (37–41). Thus, we examined whether the activation of TLR2 by the purified agonist Pam3CSK4 (referred to as Pam3) alters the transcriptome of FLAMs in a TGFβ-dependent manner. TGFβ (+) and TGFβ (-) FLAMs were stimulated with Pam3 for 18 hours, then RNA sequencing and differential expression analysis was used to identify changes in the transcriptional landscape. We identified over 700 genes that were significantly altered following Pam3 treatment of TGFβ (+) FLAMs compared to untreated TGFβ (+) FLAMs (Supplementary Figure 2A and GSE276577 for normalized counts). We were next curious which pathways were enriched among differentially expressed genes in TGFβ (+) FLAMs compared to TGFβ (-) FLAMs following Pam3 treatment to identify TGFβ-dependent, and perhaps, AM-specific immune signaling (Figure 2A). Using GSEA, we found an unexpected enrichment in IFNα response in the TGFβ (+) FLAMs (Figure 2B). When we examined the entire IFNα hallmark pathway, which encompasses the general type I IFN response, across all conditions we only observed robust induction of IFN-related genes in Pam3 treated TGFβ (+) FLAMs (Figure 2C). This finding suggests that TGFβ skews macrophage responses to drive the activation of type I IFN pathways following Pam3 stimulation.

**Figure 2.**
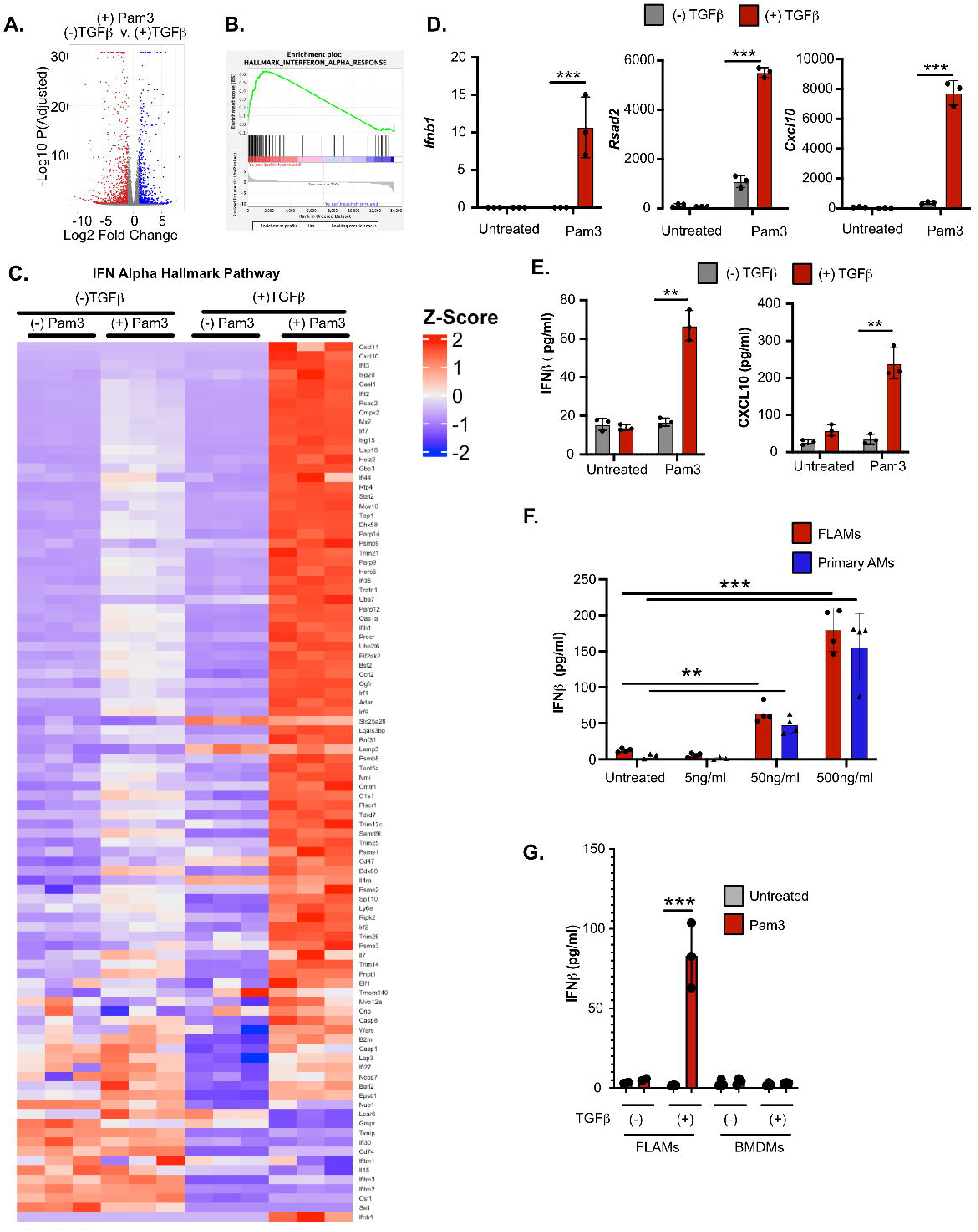
TGFβ mediates type I IFN responses during TLR2 activation. **(A)** TGFβ (+) and TGFβ (-) FLAMs were stimulated with 50 ng/ml Pam3 and 18 hours later differentially expressed genes were identified between TGFβ (+) and TGFβ (-) FLAMs left untreated or treated with Pam3 treated for 18 hours. Red points represent significantly under-expressed genes (adjusted p-value < 0.05) and blue points represent significantly overexpressed genes (adjusted p-value < 0.05) between TGFβ (+) and TGFβ (-) FLAMs. **(B)** Leading edge analysis of differentially expressed genes identified the IFN-alpha response hallmark pathway comparing Pam3 activation in TGFβ (+) and TGFβ (-) FLAMs. **(C)** Expression of genes representing the IFNα hallmark pathway between TGFβ (+) and TGFβ (-) FLAMs that have or have not been treated with Pam3. Each column of 3 biological replicates represents an experimental condition from one experiment. **(D)** Normalized read counts from *Ifnb*, *Rsad2* and *Cxcl10* from Pam3 RNA sequencing experiment. *** adjusted p-value <0.001 using DeSeq2 analysis. **(E)** TGFβ (+) and TGFβ (-) FLAMs were stimulated with Pam3 for 24 hrs. Supernatants were collected for quantification of IFNβ and CXCL10 by ELISA **(F)** TGFβ (+) FLAMs or primary AMs from C57BL6/J mice were stimulated with the indicated concentrations of Pam3 and IFNβ was quantified by ELISA the following day. Shown is one representative experiment of three each containing 3 replicates per experiment. **p<0.01 ***p<.001 by one-way ANOVA with a Tukey post-hoc test for multiple comparisons. **(G)** FLAMs or BMDMs grown with or without TGFβ were stimulated with Pam3 for 24 hours and IFNβ was quantified by ELISA. Shown is one representative experiment of two containing three replicates per group.

To further understand the functional outcomes of TLR2 dimerization in TGFβ (+) FLAMs, we next directly examined the normalized reads of IFN related genes. Our transcriptome data showed no expression of any *Ifna* gene or *Ifne* and *Ifng*, but we did observe increased expression of *Ifnb1* and the interferon-stimulated genes (ISGs) *Cxcl10* and *Rsad2* (Figure 2D), suggesting IFNβ is driving the gene signature. We observed similar baseline expression of *Ifnb1*, *Cxcl10* and *Rsad2* between conditions; however, TGFβ (+) FLAMs induced significantly higher expression of all three genes following Pam3 treatment. To corroborate the RNA sequencing results, we compared the secretion of cytokines in resting and Pam3 treated TGFβ (+) and TGFβ (-) FLAMs using an ELISA (Figure 2E). In agreement with our transcriptional results, we observed a significant increase in IFNβ and CXCL10 secretion from Pam3 treated TGFβ (+) FLAMs compared to TGFβ (-) FLAMs. In line with our Pam3-simulated TGFβ (+) FLAMs, we observed that two additional agonists that mimic bacterial or fungal infection, peptidoglycan and zymosan respectively, resulted in a significant increase in CXCL10 production in TGFβ (+) FLAMs compared to TGFβ (-) FLAMs (Supplemental Figure 2B). In contrast, we observed no significant CXCL10 secretion following activation with depleted zymosan or curdlan (Supplemental Figure 2B), which are pure Dectin-1 ligands (42).

We next confirmed this phenotype occurs in primary murine AMs by isolating cells from the lungs and treating them with increasing Pam3 concentrations, subsequently examining the production of IFNβ by ELISA (Figure 2F). In line with our results in FLAMs, we observed a significant, dose dependent increase in IFNβ in AMs following treatment with Pam3. To determine if this innate response was AM specific, we next examined primary bone marrow-derived macrophages (BMDMs). FLAMs or BMDMs were grown with or without TGFβ then stimulated with Pam3 and the following day, IFNβ was quantified by ELISA (Figure 2G). We only observed increased IFNβ secretion from FLAMs grown in TGFβ. These data suggest that the effects of TGFβ on TLR2 signaling may be specific to tissue-resident macrophages.

### TLR2 signaling in TGF**β** (+) FLAMs induces an RLR gene signature and type I IFN production is dependent on MAVS, IRF3, and IRF7

Since our results suggested that TLR2 agonists drive an increased type I IFN response in TGFβ (+) FLAMs, we next directly tested the role of TLR2 signaling in the induction of the type I IFN response using *Tlr2^(-/-)^* FLAMs. Wild type and *Tlr2^(-/-)^* TGFβ (+) FLAMs were stimulated with Pam3 and the IFNβ protein concentrations were quantified in the supernatants by ELISA the following day. While wild type TGFβ (+) FLAMs robustly induced IFNβ, this was lost in *Tlr2^(-/-)^* FLAMs (Figure 3A). This result suggests that TGFβ signaling in FLAMs drives a unique response to TLR2 activation that results in the production of type I IFN.

**Figure 3.**
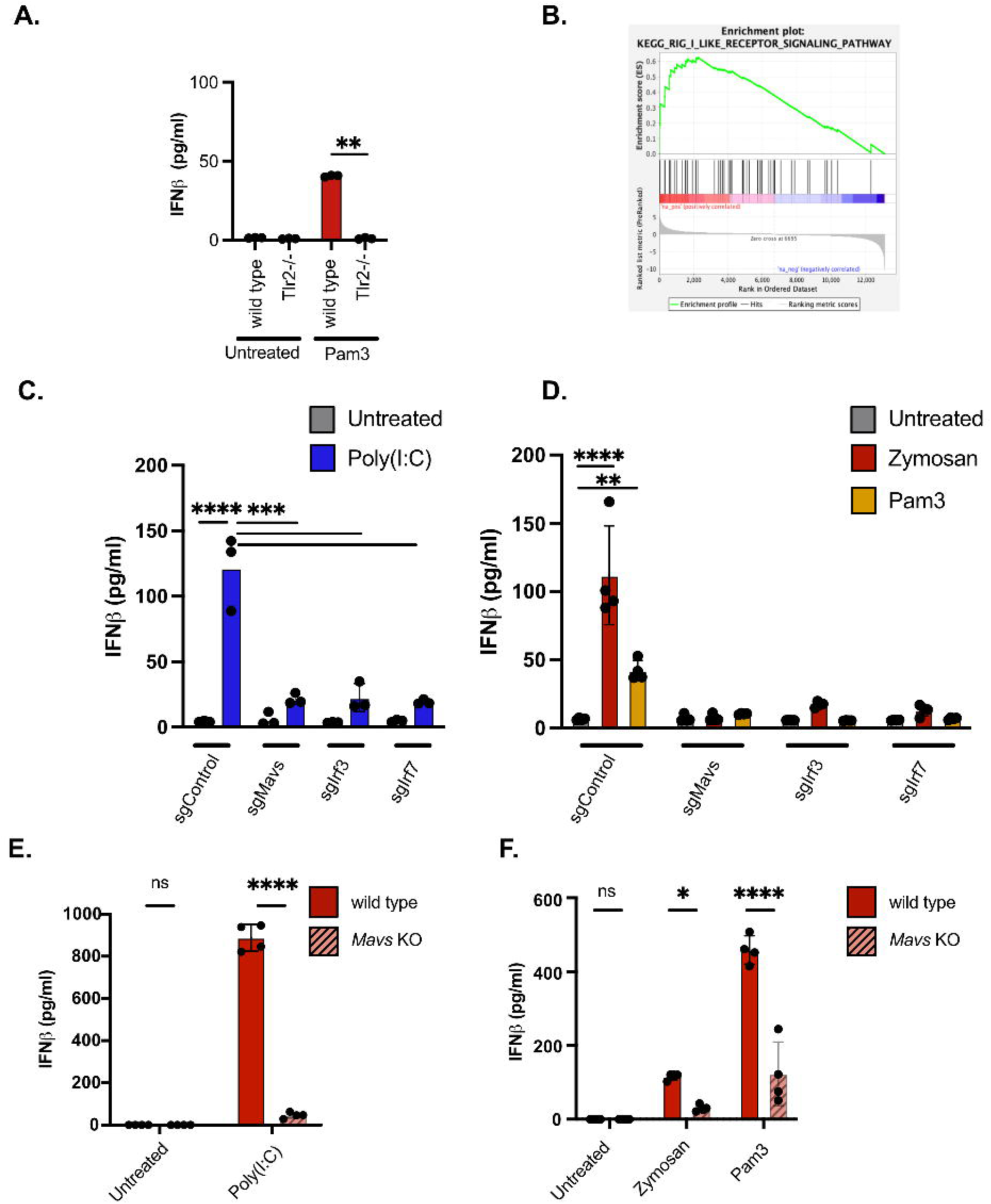
MAVS contributes to TGFβ-dependent type I IFN responses. **(A)** Wild type FLAMs or *Tlr2^(^*^-/-)^ FLAMs, were stimulated with Pam3 for 24hrs. Secreted IFNβ was quantified by ELISA. Each point represents data from a single well from one representative experiment of three. ****p<0.0001 ** p<0.01 by one-way ANOVA with a Tukey post-hoc test for multiple comparisons. **(B)** Leading edge analysis of differentially expressed genes identified RLR signaling comparing Pam3 activation in TGFβ (+) and TGFβ (-) FLAMs. **(C)** Wild type, sgMAVS, sgIRF3, and sgIRF7 FLAMs were stimulated with Poly(I:C) for 24hrs. Secreted IFNβ was quantified by ELISA. Shown is a representative experiment of three with at least 3 samples per group. **(D)** Wild type, sgMAVS, sgIRF3, and sgIRF7 FLAMs were stimulated with Pam3 or Zymosan (1μg/ml) for 24hrs. Secreted IFNβ was quantified by ELISA. Shown is a representative experiment of three with at least 3 samples per group. **(E)** Wild type or *Mavs* KO FLAMs were stimulated with Poly(I:C) for 24hrs. Secreted IFNβ was quantified by ELISA. **(F)** Wild type or *Mavs* KO TGFb (+) FLAMs were stimulated with Pam3 or Zymosan (1μg/ml) for 24hrs. Secreted IFNβ was quantified by ELISA. Each point represents data from a single well from one representative experiment of four. *** p<0.001, ** p<0.01 by one-way ANOVA with a Tukey post-hoc test for multiple comparisons.

We next wanted to better understand the pathways driving the TLR2-dependent type I IFN response in TGFβ (+) FLAMs. Examining our transcriptional analysis in Pam3 treated TGFβ (+) and TGFβ (-) FLAMs, we observed a signature GSEA enrichment for RLR signaling (Figure 3B). RLR signaling converges on the mitochondrial antiviral-signaling (MAVS) protein which triggers the activation of the transcription factors interferon regulatory factors 3 and 7 (IRF3/IRF7) to mediate the transcription of IFNβ (43, 44). To test the role of these genes in controlling TGFβ-dependent type I IFN responses after Pam3 stimulation, we used our previously described CRISPR-Cas9 editing approaches in FLAMs to target *Mavs*, *Irf3* and *Irf7* with individual sgRNAs (32).

We first confirmed the functional effects of editing by stimulating TGFβ (+) control, sgMAVS, sgIRF3, and sgIRF7 FLAMs with Poly(I:C) a known activator of MAVS-dependent RLR signaling and measured IFNβ production by ELISA (45). We observed a significant decrease in IFNβ for all three genes targeted suggesting functional defects in RLR signaling (Figure 3C). We next tested the response of these cells to Pam3 and zymosan. The following day secreted IFNβ was quantified by ELISA. Similar to our previous findings, we observed that wild type FLAMs induced IFNβ in response to both Pam3 and zymosan (Figure 3D). Surprisingly, we found IFNβ secretion after both Pam3 and zymosan treatment was significantly reduced from the sgMAVS, sgIRF3, and sgIRF7 FLAMs (Figure 3D). These data suggest that TGFβ-dependent, TLR2-mediated type I IFN responses are controlled by MAVS and IRF3/IRF7. To account for possible effects of lentiviral transduction, we next used Cas9 mediated editing with ribonuclear protein (RNP) to generate *Mavs*-deficient FLAMs. These *Mavs*-deficient cells did not produce IFNβ in response to Poly(I:C), suggesting functional knockout of MAVS (Figure 3E). In agreement with our lentiviral delivered sgRNA studies, we found that genetic loss of *Mavs* results in a reduction in IFNβ following both Pam3 and zymosan stimulation (Figure 3F). Thus, TGFβ-dependent, TLR2-mediated type I IFN responses are controlled by MAVS-dependent signaling.

### Mitochondrial oxidative stress contributes to TLR2-dependent IFNβ production

Given MAVS was required for the type I IFN response, we next wanted to understand the mechanisms driving MAVS activation and signaling. Our transcriptomics suggest that TGFβ drives lipid metabolism, and the cellular hubs of lipid metabolism, peroxisomes and mitochondria, are known to activate type I IFN responses (46, 47). Thus, we hypothesized that TGFβ-dependent changes to peroxisomes and mitochondria levels and/or function may condition FLAMs to drive MAVS-dependent IFNβ secretion following TLR2 activation. To test this hypothesis, we first examined peroxisomes and mitochondria in resting cells by immunofluorescent staining coupled with measuring the mean fluorescence intensity (MFI) in wild type TGFβ (-) FLAMs in addition to wild type and *Mavs* KO TGFβ (+) FLAMs. To visualize the presence of peroxisomes, we examined both PMP70, a peroxisome membrane associated protein and catalase, a luminal peroxisome enzyme (Figure 4A). We observed a significant increase in the MFI of both markers in wild type TGFβ (+) FLAMs compared to TGFβ (-) FLAMs (Figure 4B). We also noted that *Mavs* KO FLAMs resulted in a significant decrease in the MFI of peroxisomal markers, which was more like the TGFβ (-) FLAMs.

**Figure 4.**
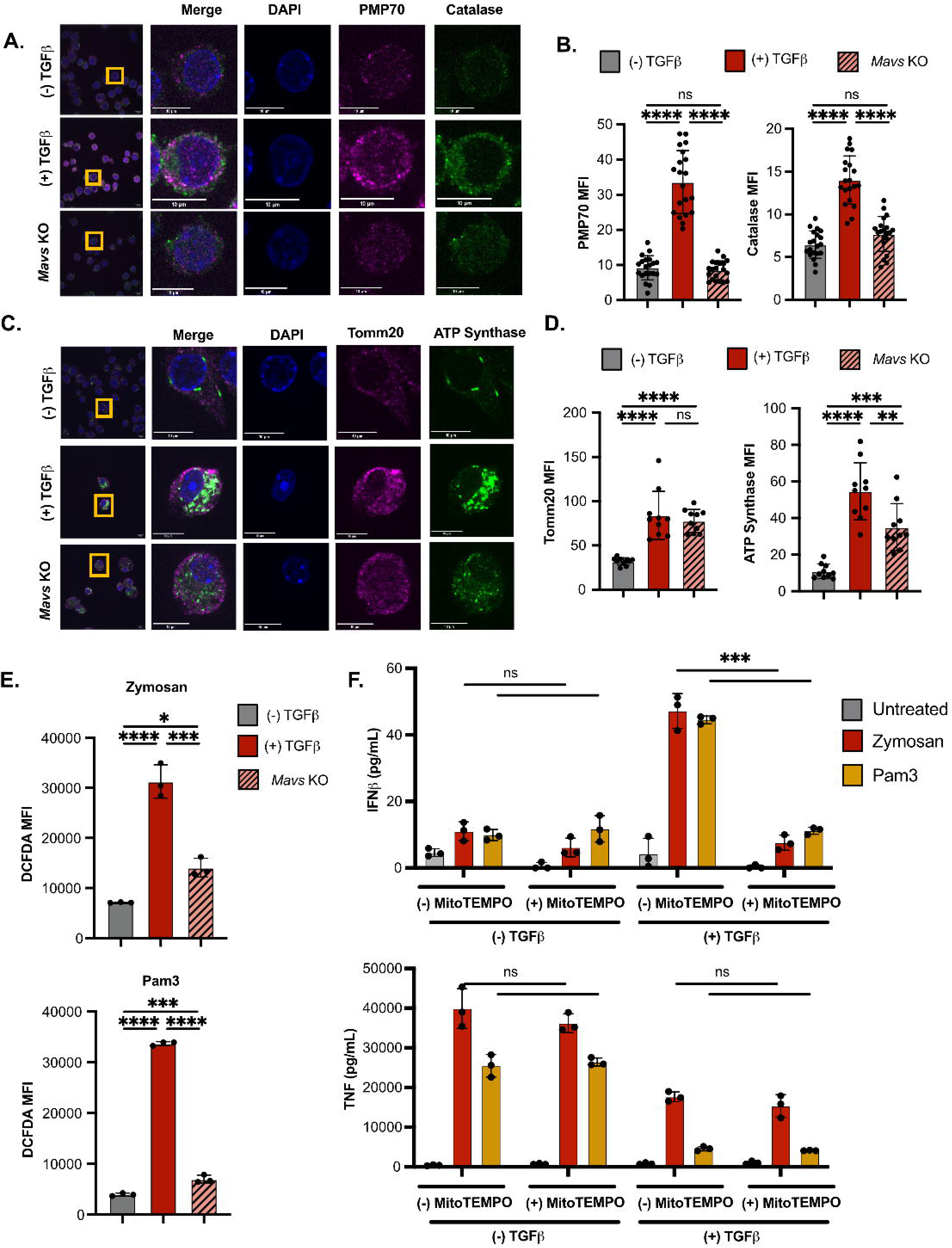
Mitochondrial ROS contributes to TGFβ-dependent IFN responses. **(A)** Wild type TGFβ (-) FLAMs and wild type or *Mavs* KO TGFβ (+) FLAMs were stained for the peroxisomal markers PMP70 and Catalase in addition to cellular DNA with DAPI. Shown are representative images from at least 3 independent experiments. The far-left column images were taken with a 100X oil objective and insets images were cropped to a single cell and have a scale bar which is 10 μm in size. **(B)** The mean fluorescence intensity (MFI) of each peroxisomal marker in (A) was quantified. Each individual data point represents a single cell from a representative experiment. **(C)** Wild type TGFβ (-) FLAMs and wild type or *Mavs* KO TGFβ (+) FLAMs were stained for the mitochondrial markers Tomm20 and ATP synthase in addition to cellular DNA with DAPI. Shown are representative images from at least 3 independent experiments. The far-left column images were taken with a 100X oil objective and insets images were cropped to a single cell and have a scale bar which is 10 μm in size. (**D**) The mean fluorescence intensity (MFI) of each mitochondrial marker in (C) was quantified. Each individual data point represents a single cell from a representative experiment. **(E)** Wild type TGFβ (-) FLAMs and wild type or *Mavs* KO TGFβ (+) FLAMs were incubated with DCFDA-H2DCFDA then left untreated or stimulated with zymosan for four hours or Pam3 for one hour followed by flow cytometry analysis. Shown is the MFI for DCFDA-H2DCFDA staining of live cells. Each point represents data from a single well from one representative experiment of two experiments with Pam3 and four experiments with zymosan. **(F)** Wild type TGFβ (-) and TGFβ (+) FLAMs were left untreated or activated with zymosan or Pam3 in the presence of absence of MitoTEMPO. 18 hours later secreted IFNβ and TNFα were quantified by ELISA. Each point represents data from a single well from one representative experiment of four. *** p<0.001, ** p<0.01 by one-way ANOVA with a Tukey post-hoc test for multiple comparisons.

We next repeated this experiment using the mitochondrial markers Tomm20 and ATP synthase (Figure 4C). We found that wild type TGFβ (+) FLAMs had a two-fold increase in the Tomm20 MFI and a five-fold increase in the MFI of ATP synthase compared to TGFβ (-) FLAMs (Figure 4D). In contrast, we found that *Mavs* KO TGFβ (+) FLAMs showed a similar increased staining of Tomm20 compared to wild type TGFβ (+) FLAMs but showed a reduction in ATP synthase staining. These data suggest that peroxisome and mitochondrial numbers are increased in TGFβ (+) FLAMs prior to activation and these organelles may be altered by the loss of MAVS.

Since lipid metabolism is a robust source of reactive oxygen species (ROS), we next hypothesized that increased peroxisomes and mitochondria levels would result in increased ROS, which could enhance MAVS-dependent signaling (48, 49). To begin to test this hypothesis, we initially quantified total ROS in cells using the dye DCFDA-H2DCFDA in wild type TGFβ (-) FLAMs in addition to wild type and *Mavs* KO TGFβ (+) FLAMs following activation with Pam3 or zymosan (Figure 4E). We observed an approximate 3-fold increase in the DCFDA-H2DCFDA MFI in wild type TGFβ (+) FLAMs following activation with either Pam3 or zymosan compared to wild type TGFβ (-) FLAMs and *Mavs* KO TGFβ (+) FLAMs. These data suggest a significant increase of ROS in TGFβ (+) FLAMs following TLR2 activation. To understand the functional consequences of increased ROS we next blocked ROS with treatment of the mitochondrial specific antioxidant MitoTEMPO (50). Both wild type TGFβ (+) and TGFβ (-) FLAMs were stimulated with Pam3 or zymosan in the presence or absence of MitoTEMPO, then the release of IFNβ and TNF were quantified by ELISA (Figure 4F). We observed that while MitoTEMPO had no effect on IFNβ release in TGFβ (-) FLAMs, there was a significant ∼4-fold decrease in IFNβ in mitoTEMPO treated TGFβ (+) FLAMs activated with both Pam3 and Zymosan. In contrast, we observed higher overall levels of TNF release from TGFβ (-) FLAMs compared to TGFβ (+) FLAMs following Pam3 or zymosan activation, but we observed no effect of MitoTEMPO on TNFα release. Taken together, ROS production, likely a result of increase fatty acid metabolism due to elevated peroxisome and mitochondria numbers, appears to be increased in TGFβ (+) FLAMs resulting in increased mitochondrial oxidative stress following TLR2 activation that contributes to the TGFβ-dependent IFN responses in the TGFβ (+) FLAMs.

## DISCUSSION

TGFβ signaling is essential for alveolar macrophage (AM) development and homeostasis in the lung environment (6, 14), but how TGFβ signaling regulates the response of AMs to external stimuli remains unclear. Here, we used an *ex vivo* model of AMs, known as FLAMs, to dissect transcriptional changes in AM-like cells that are mediated by TGFβ. Importantly, our results from this *ex vivo* FLAM model align with published results from primary AMs in the lungs, suggesting that FLAMs recapitulate many aspects of important biology observed in the lungs (6). Leveraging this powerful FLAM model, we have now found that while TGFβ restrains a subset of inflammatory pathways, TGFβ primes AMs for mitochondrial oxidative stress and a type I IFN response following treatment with TLR2 agonists. Surprisingly, these TGFβ regulated, FLAM-specific responses were regulated by MAVS- and IRF3/7-dependent signaling events downstream of TLR2 activation. These results suggest that distinct innate immune signaling networks in AMs are regulated by the tissue environment and directly alter the inflammatory response following the activation of TLR2.

While our findings demonstrate an unexpected link between TLR2, MAVS, and type I IFN production in AMs, the cell biology behind how TLR2 activates type I IFN secretion remains an open question. Several pattern recognition receptors (PRRs), including TLR3, TLR7 and TLR9 activate type I IFNs through the activation of IRF3 or IRF7, but these PRRs are localized to the endosome and generally respond to viral ligands (51, 52). In contrast, TLR2 is present on both the surface and in the endosome, similar to TLR4 (53). Previous studies showed that TLR4 signaling through the plasma membrane drives Myd88-dependent NFκβ activation, while signaling through the endosome activates a TRIF-dependent type I IFN response (54). Whether the localization of TLR2 drives the type I IFN response in TGFβ cultured AMs and how the adaptors, MyD88 and TRIF, contribute to this response will need to be determined.

While several previous studies suggest TLR2 can activate type I IFNs, the ligands and cell types capable of this response remain controversial (55–58). For example, Barbalat et al. showed BMDMs can make type I IFN in response to viral ligands but not bacterial ligands, while Dietrich et al. showed BMDMs can make type I IFN following activation with bacterial ligands (56, 57). Our data support the role of bacterial and fungal TLR2 ligands in activating a type I IFN response in AMs that is dependent on TGFβ signaling. FLAMs grown in the absence of TGFβ did not robustly induce type I IFNs following TLR2 activation. Our genetic studies found that IRF3, IRF7, and MAVS were all required for the TLR2-dependent type I IFN response. This suggests TLR2-mediated type I IFN may activate parallel pathways, one dependent on direct signaling through MyD88/TRIF, and a second dependent on the cytosolic nucleotide sensing pathways dependent on MAVS. Our data suggest a model where TGFβ primes AMs to enhance the activation of MAVS-dependent type I IFN production.

The mechanisms underlying TGFβ priming of the type I IFN response in AMs remain unknown. TGFβ is known to activate PPARγ and fatty acid oxidation (6), which we confirmed through our transcriptional analysis. Previous studies have linked cellular metabolism and type I IFN production. Both cholesterol biosynthesis and glycolysis byproducts, such as lactate, are known to regulate the magnitude of the type I IFN response in BMDMs (59, 60). Our microscopy results and transcriptional analysis showed increased fatty acid oxidation and mitochondrial function in TGFβ (+) FLAMs, which may be an upstream driver of subsequent type I IFN responses following TLR2 activation. Since we observed increased activation of MAVS-dependent type I IFN production following TLR2 stimulation, this suggested the possibility of endogenous cellular ligands such as mitochondrial DNA amplifying the TLR2 response in AMs (46). Our results with MitoTEMPO further point towards dysfunctional mitochondria as the source of ligands for activating MAVS. Yet more detailed kinetic studies of mitochondrial dynamics and cellular metabolism are warranted, as their contribution to the production of IFNβ remains unknown. Future studies will be needed to dissect the role of fatty acid oxidation, oxidative respiration, and mitochondrial damage in driving TLR2-mediated TGFβ-dependent type I IFN responses in AMs.

Our finding that AMs are uniquely programmed by TGFβ to drive a type I IFN response suggests that these specialized resident macrophages differentially activate their inflammatory profiles in the lung environment compared to other macrophages. Understanding the consequences of a type I IFN-skewed response in the lungs is an important line of research for future studies. Type I IFNs are known to be potent regulators of antiviral immunity, suggesting the host response in the lungs is particularly tuned to respond to invading viral pathogens (44). However, type I IFNs also play a key role in controlling respiratory fungal pathogens like *Aspergillus fumigatus* (61). In several disease states however, including Systemic Lupus Erythematosus and tuberculosis, elevated type I IFNs are associated with worse disease, and blocking type I IFN has been shown to improve clinical outcomes (62–64). Our data support the role of type I IFNs as a key initial response to invading pathogens in the lungs and more broadly suggests the balance of type I IFNs can mediate protective or pathologic host responses.

Interestingly, TGFβ is produced in an inactive form by AMs in the lungs and processed into an active form by integrins on lung epithelial cells which then signal back to AMs to maintain their function (6, 18, 19). This interconnected signaling ensures that AMs are properly tuned to the airspace and suggests the lung environment is an important mediator of the enhanced type I response observed in AMs. Better understanding the underlying mechanisms driving TGFβ-dependent type I IFN may enable the development of therapeutics that modulate the balance of type I IFNs more effectively in the lungs to control infections and prevent autoinflammatory diseases.

## MATERIAL & METHODS

### Animals

Experimental protocols were approved by the Institutional Animal Care and Use Committees at Michigan State University (animal use form [AUF] no. PROTO202200127) and Dartmouth College (protocol #00002168). All protocols were strictly adhered to throughout the entire study. Six- to 8-wk-old C57BL/6J mice (catalog no. 000664), *Tlr2^(-/-)^* mice (catalog no. 004650) and Cas9^(+)^ mice (catalog no. 026179) were obtained from The Jackson Laboratory (Bar Harbor, ME). Mice were given free access to food and water under controlled conditions (humidity, 40–55%; lighting, 12-hour light/12-hour dark cycles; and temperature, 24 ± 2°C), as described previously. (32). Pregnant dams at 8–10 week of age and 14–18 gestational days were euthanized to obtain murine fetuses to generated FLAMs. Primary AMs and BMDMs were isolated from male and female mice >10 week of age.

### FLAM cell culture

Wild type and *Tlr2^(-/-)^* FLAMs were isolated as previously described (32) cultured in complete RPMI (Thermo Fisher Scientific) containing 10% FBS (R&D Systems), 1% penicillin-streptomycin (Thermo Fisher Scientific), 30 ng/ml recombinant mouse GM-CSF (PeproTech), and 20 ng/ml recombinant human TGFβ1 (PeproTech) included where indicated. Media were refreshed every 2–3 d. When cells reached 70–90% confluency, they were lifted by incubating for 10 min with 37°C PBS containing 10 mM EDTA, followed by gentle scraping.

### AM and BMDM isolation and culture

Mice were euthanized by CO_2_ exposure followed by exsanguination via the inferior vena cava. Lungs were lavaged as previously described (32). Cells were then resuspended in RPMI 1640 media containing 30 ng/ml GM-CSF (PeproTech) and 20 ng/ml recombinant human TGFβ1 (PeproTech) and plated in untreated 48- or 24-well plates. AMs were lifted from plates using Accutase (BioLegend) and seeded for experiments.

For BMDMs femurs were isolated and bone marrow was harvested following centrifugation of bones cut on one side. Bone marrow was then cultured in complete RPMI 1640 media containing 30ng/ml M-CSF (PeproTech) for 7 days in untreated 15 cm dishes. Cells were then split for experiments and treated with or without 20 ng/ml recombinant human TGFβ1 (PeproTech) prior to activation.

### Flow Cytometry

To quantify surface expression of AM markers, cells were lifted by gentle scrapping, washed with PBS, and stained with SiglecF-Bv421 (Biolegend, Cat no. 155509) CD11b-APC (Biolegend, Cat no. 101212) CD14-PE-Cy7 (Biolegend, Cat no. 123316) CD11a-PE (Biolegend, Cat no. 153103) (all diluted 1:400 in PBS). Cells were then washed 3 times in PBS and fixed with 1% formaldehyde (J.T. Baker, Cat no. JTB-2106-01) in PBS. The same gating strategy was used for all experiments and consists of live cells (Fixable Live/Dead Aqua Biolegend), forward scatter and side scatter followed by doublet exclusion. Flow cytometry was performed on a BD LSR II or an Attune CytPix at the MSU Flow Cytometry Core, and data were analyzed using FlowJo (Version 10.8.1).

### TLR2 activation

Cells were seeded in 24-well or 48-well treated culture plates cells/well and allowed to settle overnight. Cells were treated with Pam3CSK4 25-200ng/ml (Invivogen, Cat no. tlrl-pms), peptidoglycan from *S. aureus* at 50μg/ml (Invivogen, cat no. tlrl-pgns2), zymosan at 25-50μg/ml or indicated concentrations (Invivogen, Cat no. tlrl-zyn), Zymosan Depleted at 50μg/ml (Invivogen, Cat no. tlrl-zyd), Curdlan at 50μg/ml (Invivogen, Cat no. tlrl-curd). As a control of direct RLR activation was mediated by treatment with 20μg/mL Poly(I:C) (Invivogen, Cat no. tlrl-pic-5) complexed with Lyovec for transfection prior to stimulation.

### Cytokine analysis

Where indicated secreted CXCL10 and TNF were quantified using the R&D Duoset kit (R&D Sciences) per manufacturer’s instructions. Secreted IFNβ was quantified with the LumiKine Xpress mIFN-B 2.0 kit (Invivogen, catalog no luex-mIFNβv2) per manufacturer’s instructions. Luminescent signal was detected on a Spark^®^ multimode microplate reader (Tecan).

### Lactate Analysis

Intracellular levels of lactate were determined using an L-lactate assay kit (Millipore Sigma, Cat. # MAK329) according to manufacturer instructions.

### Fluorescence Microscopy Staining and Imaging

FLAMs grown with or without TGFβ were seeded 18 hours before immunofluorescence staining in glass-bottom MatTek dishes (MatTek P35G-1.5-14-C) and incubated under 2mL of growth media overnight at 37°C/5% CO_2_. The following day, FLAM media was aspirated and washed with PBS then fixed with prewarmed 4% paraformaldehyde for 20 min. Following PBS washes, FLAMs were permeabilized with 0.1% Triton X-100 and blocked with 10% newborn calf serum and 0.02% NaN3. Cells were then incubated with primary antibody solution (10% newborn calf serum, 0.02% NaN3, PBS and 1° Abs) for 90 min at room temperature with the following 1° antibody concentrations: 14μg/mL PMP70:AlexaFluor647 (Invitrogen; PA1-650-A647), 10μg/mL Catalase (R&D Systems; AF3398), 4.5μg/mL Tomm20 (Abcam; AB78547), 5μg/mL ATP synthase (Invitrogen; A21351). Following PBS washes, cell were incubated with secondary antibody solution (10% newborn calf serum, 0.02% NaN3, 1x PBS, 2ug/mL DAPI and 2° Abs) for 1 hour in the dark at room temperature with the following 2° antibody concentrations: 30μg/mL rabbit anti-goat IgG:FITC (Invitrogen; 31509), 40μg/mL goat anti-rat:AlexaFluor647 (Invitrogen; A21247), 40μg/mL goat anti-mouse: fluorescein (Invitrogen; F2761). Cells were then washed with PBS and stored in the dark until imaging.

For imaging analysis, asymmetric Z-stacks (Step: 0.5μm; Below: −1.00μm; Above: 10.00μm) of FLAMs were collected using a Nikon SoRa Spinning Disk Confocal Microscope and a Plan Apochromat Lambda D 100x Oil Immersion Objective Lens, N.A. 1.45. Images were processed using FiJi ImageJ.

### ROS Quantification and MitoTEMPO treatment

ROS was quantified using DCFDA-H2DCFDA staining according to kit protocols (Abcam; ab113851). Briefly, FLAMs are incubated with 20 μM DCFDA-H2DCFDA for 30 min then were treated with 50 μM THBP as a positive control for up to 4hr at 37°C. FLAMs were stimulated with 50 μg/mL zymosan for 4 hours or 100 ng/ml Pam3CSK4 for 1 hour prior to lifting and immediate flow cytometry analysis using a Cytek Aurora in the DartLab Flow Cytometry Core. To block mitochondrial ROS, at the time of activation FLAMs were treated simultaneously with 500 μM MitoTEMPO (Sigma; SML0737) which remained in the media for the duration of the experiment. IFNβ was quantified from the supernatants by ELISA as described.

### qRT PCR

RNA from FLAMs was extracted using the Directzol RNA Extraction Kit (Zymo Research, Cat no. R2072) according to the manufacturer’s protocol. Quality was assessed using NANODROP. The One-step Syber Green RT-PCR Kit (Qiagen, Cat no. 210215) reagents were used to amplify the RNA and amplifications were monitored using the QuantStudio3 (ThermoFisher, Cat no. A28567).

*Pparg* Forward: TCGCTGATGCACTGCCTATG

*Pparg* Reverse: GAGAGGTCCACAGAGCTGATT

*Gapdh* Forward: AGGTCGGTGTGAACGGATTTG

*Gapdh* Reverse: TGTAGACCATGTAGTTGAGGTCA

### CRISPR-targeted knockouts

#### Lentiviral-mediated

Single-guide RNA (sgRNA) cloning sgOpti was a gift from Eric Lander and David Sabatini (Addgene plasmid no. 85681) (65). Individual sgRNAs were cloned as previously described (66). In short, sgRNA targeting sequences (*Irf7*: TGTGCGGCCCTTGTACATGA *Mavs*: GAGGACAAACCTCTTGTCTG *Irf3*: GGCTGGACGAGAGCCGAACG) were annealed and phosphorylated, then cloned into a dephosphorylated and BsmBI (New England Biolabs) digested sgOpti. sgRNA constructs were then packaged into lentivirus as previously described and used to transduce early passage Cas9^+^ FLAMs. Two days later, transductants were selected with puromycin. After 1 week of selection, cells were validated for SiglecF/CD14 expression and used for experimentation.

#### Ribonuclear protein-mediated

Three synthetic lyophilized sgRNAs (Synthego) per gene were pooled and prepared according to provided instructions. sgRNAs targeting *Mavs* and NLSx2-Cas9 protein were mixed with FLAMs in P2 primary cell nucleofection solution (Lonza, catalog no. V4XP-2024). Nucleofection was carried out using the 4D-Nucleofector® Core and X Unit (Lonza, catalog no. AAF-1003B and AAF-1003X).

*Mavs* guide sequences:

#1: AGGAAGCCCGCAGUCGAUCC

#2: UCUUCAAUAAUCUCCAGCGC

#3: UGCAGAUCUGUGAGCUGCCU

#### Editing efficiency by ICE analysis

Genomic DNA was isolated from control and target cells using the Qiagen DNeasy Blood and Tissue Kit (Qiagen, catalog no. 69506). Genomic DNA was quantified using a NanoDrop 1000 spectrophotometer (Thermo Scientific, catalog no. 2353-30-0010) and the edited region was amplified using polymerase chain reaction (PCR) and the samples were then sequencing using Sanger Sequencing (Genewiz) and editing efficiency was determined using ICE analysis (Synthego) (67). All cells used have an efficiency >90%.

### RNA-sequencing analysis

FLAMs with and without TGFβ were plated in 6-well plates at 1 x 10^6^ cells/well and treated with Pam3 as described above for 18 hours. We used the Direct-zol RNA Extraction Kit (Zymo Research, Cat no. R2072) to extract RNA according to the manufacturer’s protocol. Quality was assessed by the MSU Genomics Core using an Agilent 4200 TapeStation System. The Illumina Stranded mRNA Library Prep kit (Illumina, Cat no. 20040534) with IDT for Illumina RNA Unique Dual Index adapters was used for library preparation following the manufacturer’s recommendations but using half-volume reactions. Qubit™ dsDNA HS (ThermoFischer Scientific, Cat no. Q32851) and Agilent 4200 TapeStation HS DNA1000 assays (Agilent, Cat no. 5067-5584) were used to measure quality and quantity of the generated libraries. The libraries were pooled in equimolar amounts, and the Invitrogen Collibri Quantification qPCR kit (Invitrogen, Cat no. A38524100) was used to quantify the pooled library. The pool was loaded onto 2 lanes of a NovaSeq S4 flow cell, and sequencing was performed in a 2×150 bp paired end format using a NovaSeq 6000 v1.5 100-cycle reagent kit (Illumina, Cat no. 20028316). Base calling was performed with Illumina Real Time Analysis (RTA; Version 3.4.4), and the output of RTA was demultiplexed and converted to the FastQ format with Illumina Bcl2fastq (Version 2.20.0).

RNAseq analysis was completed using the MSU High Performance Computing Center (HPCC). FastQC (Version 0.11.7) was used to assess read quality. Bowtie2 (Version 2.4.1) (68) with default settings was used to map reads with the GRCm39 mouse reference genome. Aligned reads counts were assessed using FeatureCounts from the Subread package (Version 2.0.0) (69). Differential gene expression analysis was conducted using the DESeq2 package (Version 1.36.0) (70) in R (Version 4.2.1). All raw sequencing data, raw read counts, and normalized read counts are available through the NCBI Gene Expression Omnibus database (Accession: GSE276577).

Core AM upregulated signature genes were compared between FLAMs grown with and without TGFβ from our study and AMs and peritoneal macrophages from ImmGen (GSE122108 (6, 36)). Raw counts were compiled and normalized in DESeq2. A box plot was generated in GraphPad Prism using normalized counts for core AM upregulated signature genes. Gene set enrichment analysis (GSEA) was used to identify enriched pathways in the RNA-seq dataset. Genes in the indicated comparisons were ranked using DeSeq2, and the “GSEA preranked” function was used to complete functional enrichment using default settings for hallmark pathways from mice. We acknowledge our use of the gene set enrichment analysis, GSEA software, and Molecular Signature Database (MSigDB) (59) (http://www.broad.mit.edu/gsea/).

## ACKNOWLEDGMENTS

This work was supported by grants from the National Institutes of Health: R01 AI139133 (J.J.O) and R35 GM146795 (A.J.O.). A.W.R. was supported by the Dartmouth College Molecular Microbiology & Pathogenesis Program (NIH/NIAID T32 AI007519). A.P.M. was supported by the Dartmouth College Immunology Training Program (NIH/NIAID T32 AI007363). F.L. was supported by the Dartmouth College Cystic Fibrosis Training Program (NIH/NIAID T32 HL134598) and CFRDP Training Core from the Cystic Fibrosis Foundation. The Attune CytPix, located in the MSU Flow Cytometry Core Facility, is supported by the Equipment Grants Program, award #2022-70410-38419, from the U.S. Department of Agriculture (USDA), National Institute of Food and Agriculture (NIFA). DartLab, the Immune Monitoring and Flow Cytometry Shared Resource at the Dartmouth Cancer Center is partially supported with NCI Cancer Center Support Grant 5P30 CA023108-41. The Light Microscopy Facility at Dartmouth is partially supported by bioMT through NIH grant P20-GM113132, DartCF through NIH grant P30-DK117469, and by the Dartmouth Cancer Center through NCI Cancer Center Support Grant P30 CA023108. The purchase of the Nikon SoRa Spinning Disk Confocal Microscope was supported through NIH award S10 OD032310 to Dr. Yashi Ahmed (Dartmouth College). The authors wish to thank Dr. Henry Higgs, Ann Lavanway, Dr. Daniel Mielcarz, Pat Robinson, and Gary Ward for technical support at Dartmouth College.

---

**Supplemental Figure 1. (A)** Gene expression was quantified from normalized counts for key genes important in lipid metabolism, inflammation, and TGFβ signaling between TGFβ FLAMs (+) and TGFβ (-) at homeostasis. **(B)** Lactate levels produced over 18 hours were quantified from cell lysates of FLAMs grown with or without TGFβ. Shown are representative data of two experiments containing 3-5 replicates per group. Each point represents data from a single well, with the bar showing the mean ± one standard deviation. ****p<.0001 by unpaired student t-test

**Supplemental Figure 2. (A)** Differentially expressed genes were identified between TGFβ FLAMs (+) with and without Pam3 treatment for 18 hours. Red points represent under-expressed genes and blue points represent overexpressed genes following Pam3 activation. Each point represents the mean of three biological replicates from one experiment. **(B)** TGFβ (+) and TGFβ (-) FLAMs were stimulated with 50ng/ml peptidoglycan, zymosan, depleted zymosan, or curdlan for 24hrs. CXCL10 was quantified by ELISA.

