## Supplementary figures and images for "TGFβ primes alveolar-like macrophages to induce type I IFN following TLR2 activation"

### Supplemental Figure 1

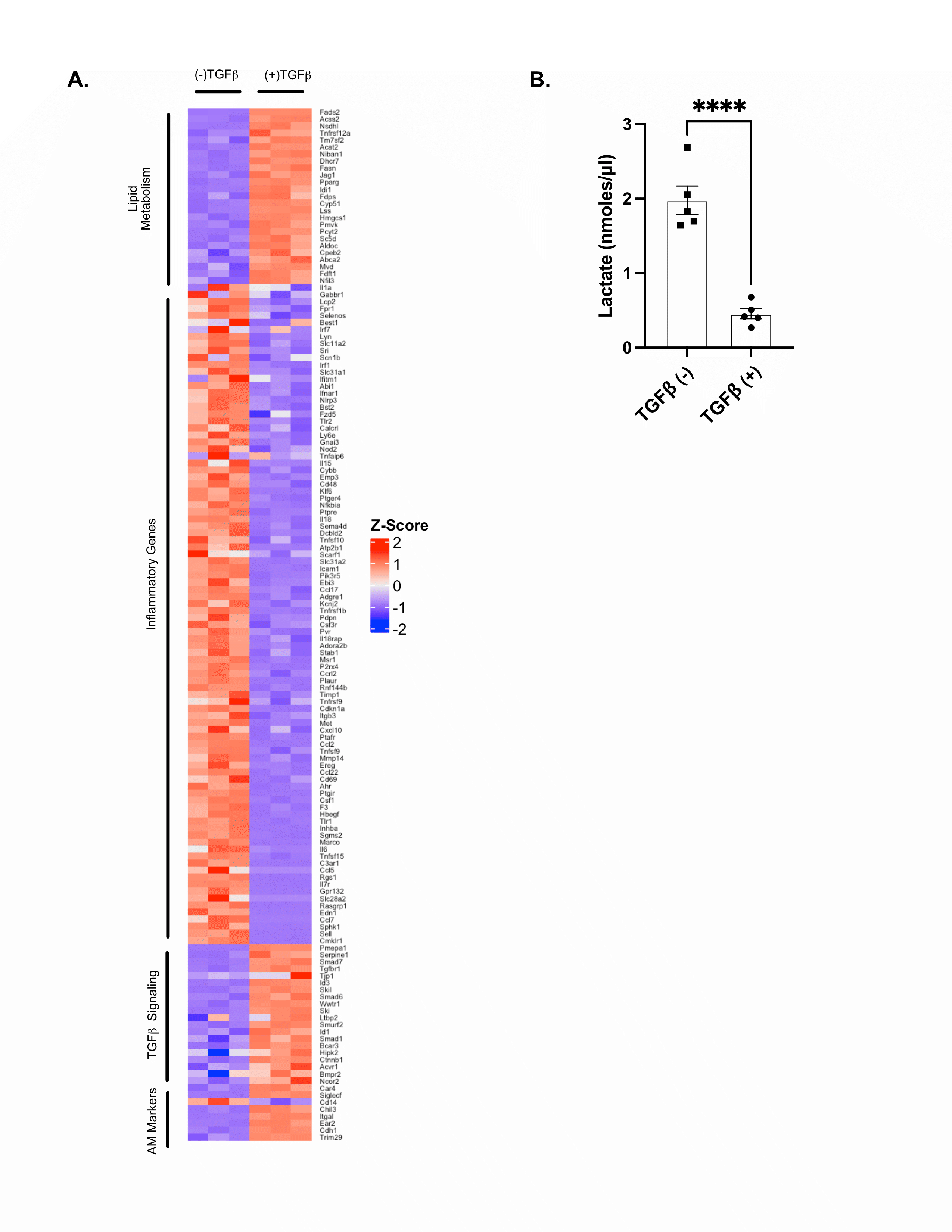

### Supplemental Figure 2

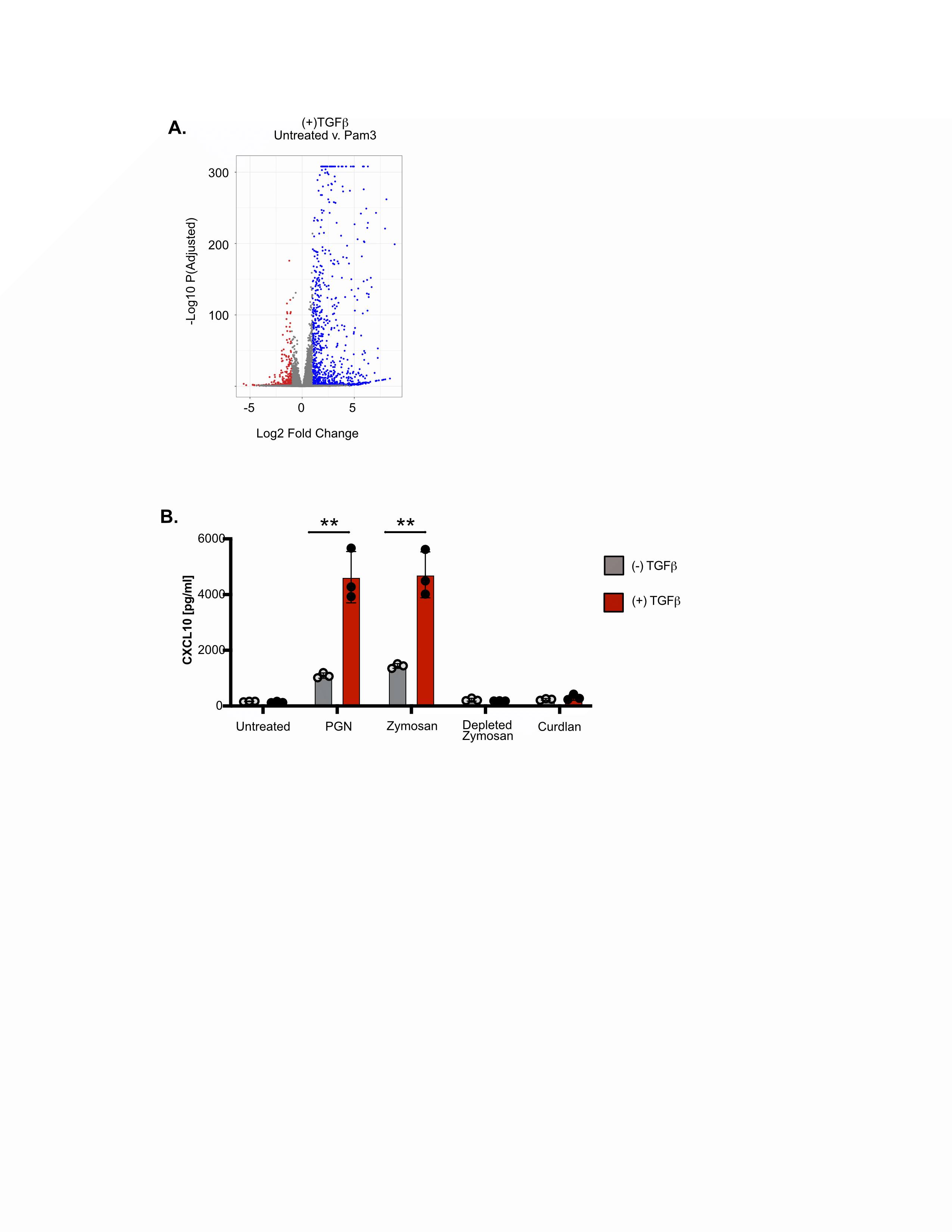
